# Splice variants of DOMINO control *Drosophila* circadian behavior and pacemaker neuron maintenance

**DOI:** 10.1101/395848

**Authors:** Zhenxing Liu, Ye Niu, Vu H. Lam, Joanna C. Chiu, Yong Zhang

## Abstract

Circadian clocks control daily rhythms in physiology. In *Drosophila*, the small ventral lateral neurons (sLN_v_s) expressing PIGMENT DISPERSING FACTOR (PDF) are the master pacemaker neurons. Despite the importance of sLN_v_s and PDF in circadian behavior, little is known about factors that control sLN_v_s maintenance and PDF accumulation. Here, we identify the *Drosophila* SWI2/SNF2 protein DOMINO (DOM) as a key regulator of circadian behavior. Depletion of DOM eliminates morning anticipation and impairs rhythmicity. Interestingly, the two splice variants of DOM, DOM-A and DOM-B have distinct circadian functions. DOM-A depletion leads to arrhythmic behavior, while DOM-B knockdown lengthens circadian period. Both DOM-A and DOM-B bind to the promotor regions of key pacemaker genes *period* and *timeless*, and regulate their protein expression. Furthermore, we identify that DOM-A is required for the maintenance of sLN_v_s and transcription of *pdf*. Lastly, constitutive activation of PDF-receptor signaling rescued the arrhythmia and period lengthening of DOM downregulation. Taken together, our findings reveal that splice variants of DOM play distinct roles in circadian rhythms through regulating abundance of pacemaker proteins and sLN_v_s maintenance.

## Introduction

Circadian clocks allow animals to anticipate daily oscillations in behavior, physiology and metabolism [1]. The core of the molecular clock is a negative transcriptional-translational feedback loop, which is evolutionarily conserved across species [2]. The fruit fly *Drosophila melanogaster* has been a powerful model in dissecting the molecular and neuronal mechanisms of circadian rhythms. In *Drosophila*, a heterodimeric complex of CLOCK (CLK) and CYCLE (CYC) activates rhythmic transcription of clock-controlled genes, including the transcriptional repressor *period* (*per*) and *timeless* (*tim*). PER and TIM dimerize and repress their own transcription by blocking CLK-CYC transactivation [3]. A number of kinases (CK1, SGG, NEMO, etc) and phosphatases (PP1, PP2A) also control the circadian clock at the post-translational level [3].

Circadian locomotor rhythms are controlled by a small set of clock neurons expressing core pacemaker proteins in the brain. In each hemisphere of the fly brain, there are ∼75 clock neurons, which are divided into clusters based on their anatomical locations and functions in circadian behavior [4]. There are three groups of dorsal neurons (DN1s, DN2s, and DN3s), the lateral dorsal neurons (LNds), the lateral posterior neurons (LPNs), the large ventral lateral neurons (lLN_v_s), and the small ventral lateral neurons (sLN_v_s). The ILN_v_s and four sLN_v_s express the neuropeptide PIGMENT DISPERSING FACTOR (PDF), while the fifth sLN_v_ is PDF negative. The PDF positive sLN_v_s are the key pacemaker neurons: they control the circadian rhythmicity under constant darkness (DD) [4]. They are also critical for generating morning anticipatory activity before lights on (i.e. dawn) under light-dark cycle (LD) [5,6]. The sLN_v_s send axonal projections towards the DN1s and DN2s, and the structural plasticity of dorsal projections has been shown to be under circadian control [7-9]. PDF positive sLN_v_s and their dorsal projections are formed in 4 hrs old first instar larvae after hatching [10].

Despite the importance of sLN_v_s in the circadian behavior, little is known about the mechanisms that control the maintenance of sLN_v_s.

PDF are the most prominent neuropeptide in the regulation of circadian behavioral rhythms [11]. PDF plays a critical role in the synchronization of different groups of clock neurons by binding to PDF receptor (PDFR), and activating cAMP signaling [12-15]. Loss of PDF or PDFR abolishes morning anticipation and significantly impairs behavioral rhythmicity in DD. Not only as a critical circadian output signal, PDF has also been shown to control the molecular clock by regulating the stability of pacemaker proteins PER and TIM recently [16,17]. PDF abundance in the sLN_v_s is regulated by CLK and CYC [18-20]. Another pacemaker protein VRILLE also promotes PDF levels [21]. Despite these studies, mechanisms regulating PDF abundance remain poorly understood.

DOM plays a critical role for incorporation and removal of the only H2A variant-H2A.V in *Drosophila* [22]. DOM is a chromatin-remodeling protein, which belongs to the SWI/SNF2 DNA-dependent ATPase family [23]. DOM is involved in oogenesis, wing development, cell viability and proliferation, neuroblast maintenance and polarity, as well as in dendrite development [23-26]. In the fly genome, two alternative splicing variants of *dom* encode DOM-A and DOM-B. Interestingly, a recent study identified that DOM-A and DOM-B play distinct roles in cell-type specific development during *Drosophila* oogenesis [24].

A few chromatin remodelers have been shown to regulate circadian photo-responses and gene expression [27-29]. However, the role played by chromatin remodeling in the control of *Drosophila* circadian clocks is still largely unknown. Here, we report the functions of DOM in the regulation of circadian rhythms. Using isoform-specific RNAi and rescue, we demonstrate distinct functions of DOM-A and DOM-B in the regulation of circadian locomotor rhythms. Depletion of DOM-A in circadian neurons leads to arrhythmic behavior and long period, while DOM-B downregulation specifically lengthens the circadian period in flies. Both DOM-A and DOM-B bind to the promoters of *per* and *tim*, and regulate the abundance of PER and TIM levels, which is critical for the control of circadian period. However, DOM-A is specifically necessary for maintenance of sLN_v_s as well as accumulation of PDF in these neurons. Indeed, activation of PDFR signaling restores the locomotor rhythms of DOM downregulation. Together, our results suggest that the two alternative spliced variants of DOM play distinct roles in circadian rhythms.

## Results

### DOM regulates circadian locomotor rhythms

Mass spectrometry (MS) label-free quantitative proteomics approach was previously performed to identify the BRAHMA (BRM) chromatin-remodeling protein complex as interactor of circadian clock proteins [30]. In the same data set, we observed that several core subunits of the ATP-dependent DOM chromatin-remodeling complex, including DOM, NIPPED-A, PONT, REPT, and Mrg15, interact with CLK in the nucleus of *Drosophila* S2 cells (Table 1). Since DOM is the ATPase subunit of the protein complex and shows significant binding to CLK (especially C-terminal FLAG-tagged CLK) according to SAINT (Significant Analysis of INTeractome) scoring [31], we decided to further investigate its role in circadian regulation. Interestingly, prior studies indicated that *dom* mRNA levels in the sLN_v_s are clearly enriched about 4.4-folds compared to other neurons [32].

**Table 1.**
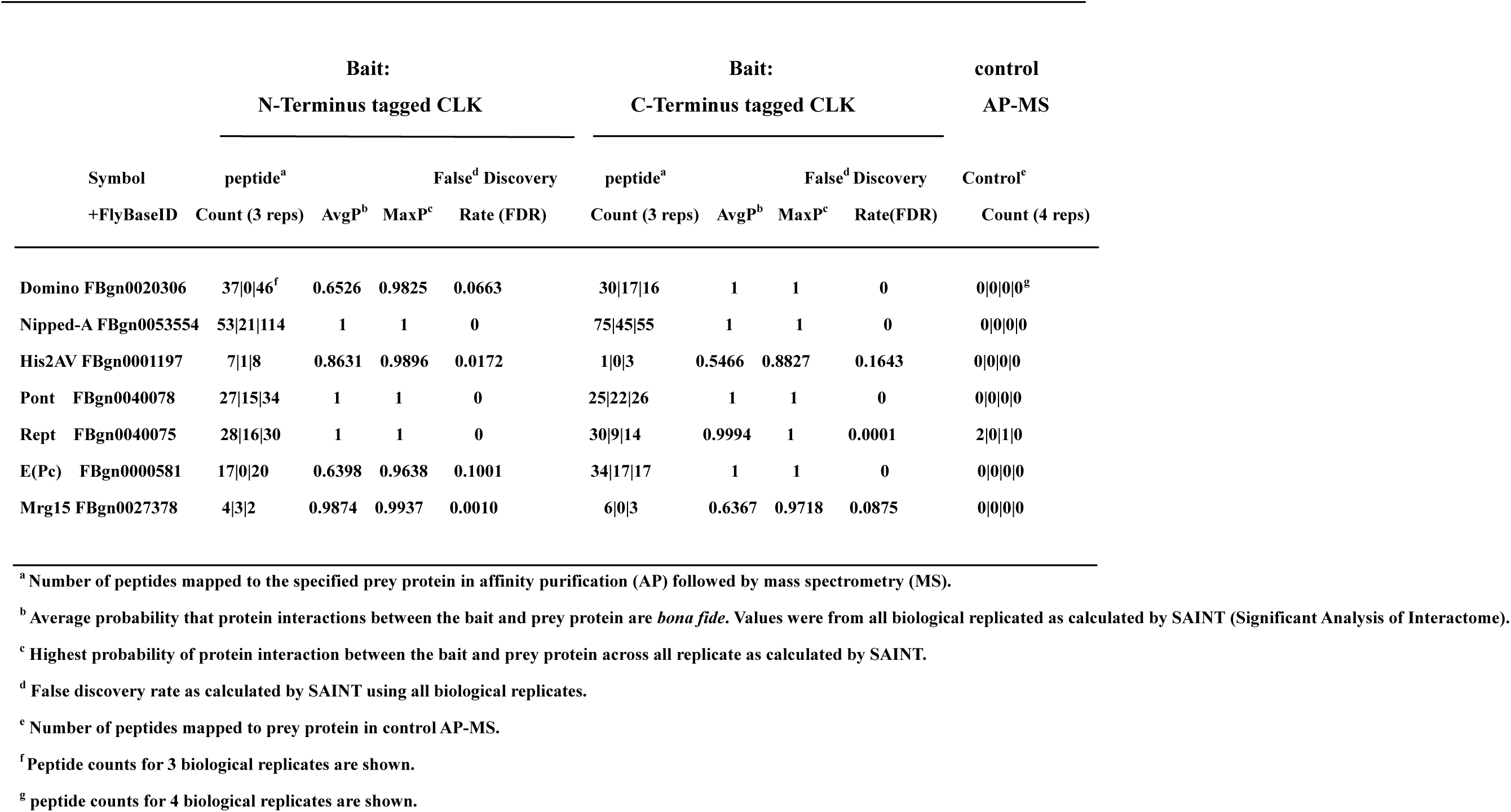
Subunits of the DOM chromatin-remodeling complex interact with CLK in the nucleus of *Drosophila* Schneider (S2) cells as detected by FLAG affinity purification followed by mass spectrometry.

Homozygous *dom* mutants are lethal, therefore we used RNAi to deplete DOM in circadian neurons. *Dicer 2* was co-expressed to enhance RNAi efficiency and has been shown to have no effects on circadian rhythms [33,34]. When we expressed *dom* dsRNAs using *tim-GAL4*, a circadian tissue-specific driver [35], most of the flies became arrhythmic, and the amplitude of rhythm was significantly reduced (Fig. 1A and Appendix Table S1). Only 31% of flies with *dom* downregulation (41% for another independent line *dom* RNAi#2) were rhythmic, and interestingly the circadian period of these lines was about 1.5 hours longer than the control. The reduced rhythmicity and amplitude were also observed when we only targeted the PDF positive LN_v_s using *pdf*-*GAL4* [11] (Fig. 1A). However, neither percentage of rhythmicity nor amplitude was affected when we targeted all circadian cells except the PDF positive LN_v_s, using a combination of *tim-GAL4* with the repressor transgene *pdf-GAL80* (Fig. 1A). However, a slight period-lengthening of activity rhythm was detected. These results indicated that DOM primarily functions in the LN_v_s to control circadian behavior, but might be needed in PDF negative circadian neurons to fine-tune circadian period length.

**Figure 1.**
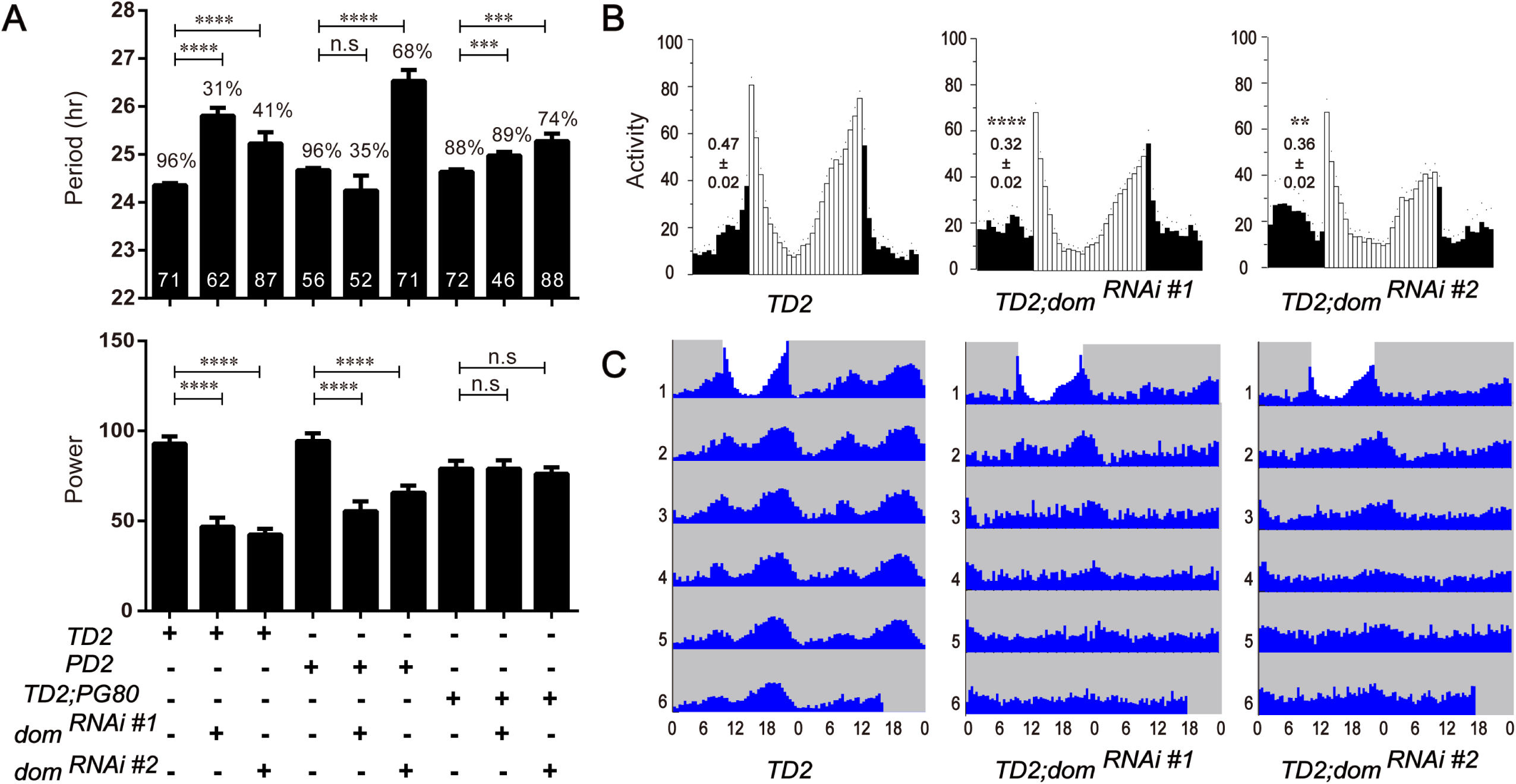
Depletion of DOM disrupted circadian rhythms. **A.** Free-running period (top panel) and power (bottom panel) of DOM depleted flies. The percentages of rhythmic flies are shown above each column. The number of tested flies is shown in each column. *dom* mRNA targeted by the two non-overlapping RNAi lines from the TRiP stock center. *dom*^*RNAi#1*^ (JF01502) is a long double-stranded RNAi line, while *dom*^*RNAi#2*^ (HMS01855) is a short hairpin RNAi line. TD2=*tim-GAL4, UAS-dcr2*, PD2= *pdf-GAL4, UAS-dcr2*, TD2; PG80= *TD2, pdf-GAL80*. *n*= 50–90; error bars represent ± SD; n.s., non-significant, **p < 0.01, ***p < 0.001, \*\*\*\**P* < 0.0001; one-way ANOVA. **B.** Average locomotor activity of flies under 3 days of 12:12 hr LD conditions. Dark activity bars represent the night, and white bars represent the day. The significant differences of the values (shown above bars) indicate morning anticipation is severely disrupted in *dom* RNAi lines. **C.** Actograms showing the average activities on the last day of LD followed by 5 days in DD. Light represents the day and gray darkness. From left to right: (Left panel) Gal4 control; (middle panel) *dom*^*RNAi#1*^ flies; and (right panel) *dom*^*RNAi#2*^ flies (knockdown of *dom* in all circadian neurons). Depletion of DOM caused arrhythmia in DD.

Because of the high arrhythmia in DD, we also examined the behavioral phenotype of *dom* RNAi flies under LD. We found that DOM knockdown abolished the morning peak of anticipatory activity in flies (Fig. 1B). Importantly, two independent *dom* RNAi lines targeting non-overlapped regions of *dom* exhibited similar phenotypes in both LD and DD (Fig. 1C and Appendix Fig S1). Thus, it is unlikely that off-target effect of *dom* RNAi cause these circadian phenotypes. Furthermore, quantitative RT-PCR results showed that *dom* expression were significantly reduced in fly heads of DOM knockdown as compared to controls (Appendix Fig S1). Together, these results suggest that DOM is important for regulation of circadian locomotor behavior.

### DOM regulates the abundance of PER and TIM

Depletion of DOM severely decreased the rhythmicity and lengthened circadian period. To understand how DOM affects circadian rhythms, we first measured the oscillations of mRNA abundance of three core pacemaker genes in fly heads. For *clk*, the abundance and oscillation of mRNA under LD and DD were not affected in *dom* RNAi flies (Fig. 2A). However, we found that the abundance of *per* and *tim* mRNA was reduced, especially at the time point of peak expression (Fig. 2A). Under DD, depletion of DOM also decreased the level of *per* and *tim* mRNA, which indicates that the effects are not due to masking effects of light (Fig. 2A). Next we used PER as a marker of molecular pacemaker and examined the oscillation of PER in pacemaker neurons across 24 hours under DD. Consistent with the changes in mRNA level, the level of PER was also significantly reduced in the sLN_v_s with DOM depletion (Fig. 2B, 2C). In other groups of circadian neurons, such as the LNds and DN1s, decreased abundance of PER was also observed (Appendix Fig S2). Taken together, our results suggest that DOM controls the abundance of PER and TIM both at mRNA and protein levels.

**Figure 2.**
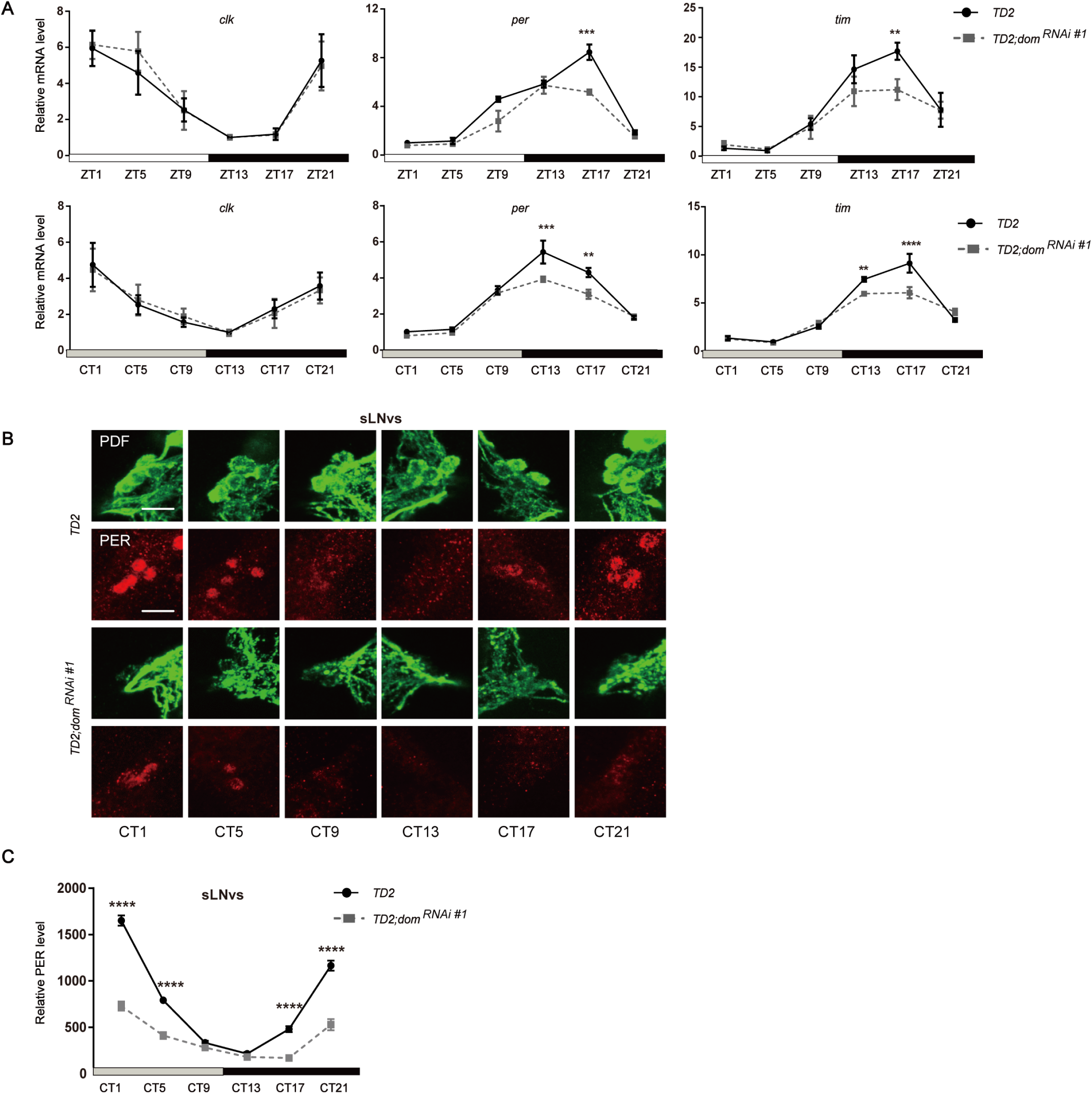
Downregulation of DOM decreases the abundance of PER and TIM. **A.** Quantitative RT-PCR showing the expression of *clk, per* and *tim*. Flies were collected at the indicated time points (ZT= Zeitgeber Time or CT= circadian time). Downregulation of *dom* decreased *per* and *tim* mRNA levels (middle and right panels), while *clk* was normal (left panel). **B.** Whole mount immunostaining showing the expression pattern of PER in sLN_v_s. Red represents PER and green is PDF. Flies were entrained for 4 days in LD and transferred to DD and dissected every 4 h on the fifth day. Downregulation of DOM decreased PER levels at CT1-5 and CT17-21. (Scale bar: 50 um.) **C.** Quantification of the staining in sLN_v_s. For each genotype, 16-20 fly brains and 60-80 neurons were used for quantification. White and black bars indicate lights-on and lights-off, respectively. Gray and black bars indicate subjective day and subjective night, respectively. Time (h) is indicated as ZT or CT where CT0 is 12 h after lights-OFF of the last LD day. Two independent experiments were done for each genotype/condition with very similar results. Error bars correspond to SEM. **p < 0.01, ***p < 0.001, ****p < 0.0001 as determined by the *t*-test.

### DOM-A but not DOM-B is regulated by CLK

There are two major splice variants of *dom*: *dom-A* and *dom-B*, which has distinct functions during Drosophila oogenesis [24]. We wondered whether these two isoforms would have different functions in circadian rhythms. We first examined the expression pattern of *dom-A* and *dom-B* in the fly head across different times of the day. Interestingly, *dom-A* exhibited a strong oscillation pattern with a trough around zeitgeber time 9 (ZT9, as ZT0 is light on and ZT12 is light off) and a peak expression near ZT21 (Appendix Fig S3A). Oscillation of *dom-A* was also observed under DD. Importantly, the oscillation of *dom-A* expression was abolished in the *Clock* null mutant *Clk*^*out*^ (Appendix Fig S3A) [36]. Together, these data indicate that *dom-A* is controlled by circadian clock. However, we did not observe a clear oscillation for *dom-B* expression (Appendix Fig S3B). In order to detect changes at protein level, we took advantage of isoform-specific antibodies to DOM-A or DOM-B (25). We confirmed the specificity of these antibodies by altering the level of DOM-A or DOM-B using RNAi and overexpression (Appendix Fig S3C-3S). Similar to mRNA level, DOM-A protein abundance was dramatically reduced in *Clk*^*out*^, however, no oscillation of DOM-A was observed in wild-type (Appendix Fig S3E,S3G). This loss of DOM-A oscillation at protein level indicates that DOM-A protein might be quite stable. The levels of DOM-B were not affected in *Clk*^*out*^ (Appendix Fig S3F,S3H).

### DOM-A and DOM-B have distinct roles in circadian regulation

Differential regulations of DOM-A and DOM-B by CLK suggest that they might have different roles in circadian rhythm. We therefore performed isoform-specific downregulation in circadian tissues using small hairpin RNA (shRNA) targeting *dom-A* or *dom-B*. These transgenic fly lines have been previously shown to specifically knockdown these alternative isoforms (Appendix Fig S1) [24]. We first quantified the efficiency of *dom-A* and *dom-B* downregulation by shRNAs in fly heads using quantitative real-time PCR. Consistent with previous report [24], we observed that each shRNA line specifically downregulated *dom-A* or *dom-B* (Appendix Fig S1B). Interestingly, when we depleted DOM-A or DOM-B in all circadian clock neurons, we observed two distinct circadian behavioral phenotypes. Most of the flies expressing *dom-A* shRNA lost locomotor rhythms, except for ∼20% that were rhythmic and showed a longer period as compared to the control (Fig. 3A, 3B and Appendix Table S1). As with depletion of all *dom* isoforms, DOM-A knockdown also blunted the morning activity peak under LD (Appendix Fig S4). However, for DOM-B knockdown, flies exhibited a lengthened period of ∼1.5 hrs (Fig. 3A, 3B), but no effect on amplitude of rhythms or morning anticipation was observed (Fig. 3B, and Appendix Fig S4). When we expressed *dom-A* or *dom-B* shRNA only in LN_v_s, or in PDF negative circadian neurons, we found similar effects on rhythmicity or period-lengthening (Fig. 3B). These results suggested that DOM-A and DOM-B are required in both PDF positive and negative circadian neurons for behavior. It is unlikely that the different circadian phenotype is due to greater efficiency of *dom-A* over *dom-B* shRNA (Appendix Fig S1B). Actually a previous study has also validated the knockdown efficiency and found that *dom-B* shRNA is in fact slightly stronger than *dom-A* in the fly nervous system [24].

**Figure 3.**
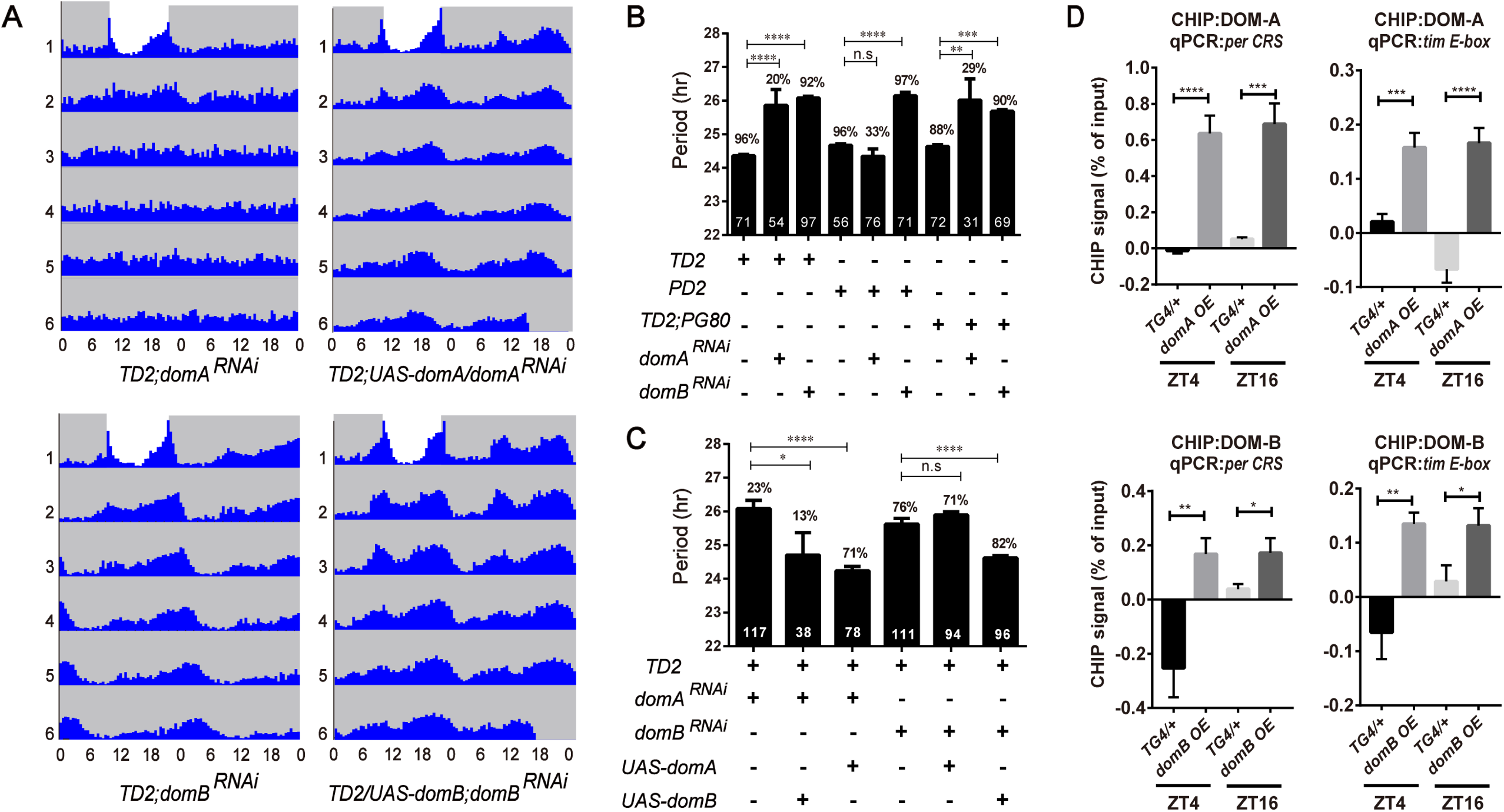
DOM-A and DOM-B have distinct functions in regulating circadian locomotor rhythms. **A.** Actograms showing the average activities on the last day of LD followed by 5 days in DD. Light represents the day and gray darkness. From top to bottom: *domA*^*RNAi*^ (Top left panel); *domA*^*RNAi*^; *UAS-domA rescue*(Top right panel); and *domB*^*RNAi*^ (Bottom left panel); *domB*^*RNAi*^; *UAS-domB rescue*(Bottom right panel). Depletion of *domA* caused arrhythmia in DD, while knocking down *domB* lengthened circadian period. *domA*^*RNAi*^ and *domB*^*RNAi*^ phenotypes can be rescued by restoring *domA* and *domB* expressing in all circadian neurons respectively. **B.** Free-running period and percentage of rhythmicity of *domA* and *domB* depleted flies. The percentages of rhythmic flies are shown above the error bar. The numbers of tested flies are shown in each column. **C.** Free-running period and percentage of rhythmicity of flies with restoring DOM-A or DOM-B in *domA* or *domB* depletion. **D.** ChIP assays detecting DOM-A and DOM-B binding more at the *per CRS* and *tim E-box* in flies expressing *domA* (BL64261) and *domB* (BL64263) in tim-expressing cells as compared to control TG4 flies. Non-specific DOM binding was detected by amplifying an intergenic region (FBgn0003638) of the *Drosophila* genome and subtracted from the signal from the *per CRS* and *tim E-box 1* signals. Results shown are from at least three biological ChIP replicates, with technical triplicates performed during qPCR for each biological replicate. Error bars represent ± SEM; n.s., non significant,\**P* < 0.05,**p < 0.01, ***p < 0.001, \*\*\*\**P* < 0.0001; one-way ANOVA.

Our isoform-specific knockdown indicated that DOM-A and DOM-B has distinct functions in circadian rhythms. Although the shRNA lines have been shown here and in a previous study [24] to specifically knockdown the *dom-A* or *dom-B* isoforms, off-target effects could still contribute to the observed phenotypes. Thus, we performed rescue experiments with UAS transgenic flies expressing either *dom-A* or *dom-B* cDNA. Circadian behavior defects were rescued when we overexpressed corresponding UAS lines in the DOM-A or DOM-B knockdown (Fig. 3A, 3C), which indicated that the isoform specific phenotype we observed was not due to off-target effects. Remarkably, we were not able to rescue the long period phenotype of DOM-B knockdown with overexpression of *dom-A* (Fig. 3C). Similarly, neither was overexpression of *dom-B* able to rescue the arrhythmic phenotype of DOM-A knockdown, although we did notice that the period-lengthening effect was partially rescued (Fig. 3C).

Since DOM is a chromatin-remodeling protein, and since we observed lower abundance of both *per* and *tim* mRNA, we therefore examined whether DOM might be bound to the promoters of *per* and *tim.* We used the DOM-A- and DOM-B-specific antibodies to perform chromatin immunoprecipitation (ChIP) at two time points: ZT4 and ZT16, close to the trough and peak time of CLK binding [37]. We observed significant enrichments of DOM-A and DOM-B binding on both *per* and *tim* promoters, compared to an intergenic region control (Fig. 3D). However, we did not observe significant difference in binding between ZT4 and ZT16 (Fig. 3D), which indicates that the binding of DOM-A and DOM-B might not be time-dependent.

### DOM-A, but not DOM-B, is required for the maintenance of sLN_v_s and PDF abundance

The blunted morning activity peak and arrhythmia in DOM and DOM-A knockdown flies suggest that there might be defects in the sLN_v_s or in PDF signaling. There are three major projections of PDF positive LN_v_s. The lLN_v_s send projections to the optic lobes and the contralateral brain hemisphere, while the sLN_v_s send projections to the DN1s and DN2s region [38]. Based on whole mount immunohistochemistry in fly brains, we did not observe obvious defects in the brain structure or projections of the lLN_v_s, however, PDF levels in the dorsal projections of the sLN_v_s were barely detectable in DOM and DOM-A knockdown flies (Fig. 4A). Absence of the dorsal projection or low PDF expression in the sLN_v_s could lead to decrease of PDF in the dorsal projection [19,39]. Our data are consistent with both possibilities. Using *pdf*-*GAL4* and a membrane tethered GFP (CD8-GFP) to label axons of sLN_v_s, we observed that the dorsal projections of sLN_v_s were clearly shortened in DOM and DOM-A but not in DOM-B downregulation (Appendix Fig S5). Furthermore, close observation of the PDF positive sLN_v_s revealed that both the number of sLN_v_s and PDF levels were reduced (Fig. 4B, 4C, and Appendix Fig S5). For each hemisphere, the average number of PDF positive sLN_v_s in DOM and DOM-A knockdown flies was approximately 2, compared to 4 in the control flies (Fig. 4C). However, with DOM-B knockdown, we did not observe any obvious loss of sLN_v_s or reduction in PDF levels, which indicates that this process is specifically controlled by DOM-A (Fig. 4B, 4C). Consistent with the ChIP results, the levels of PER and TIM were found to be significantly decreased in the sLN_v_s with knockdown of DOM-A or DOM-B (Fig. 4B, 4C). This may also explain why depletion of DOM-A or DOM-B had an effect on period-lengthening.

**Figure 4.**
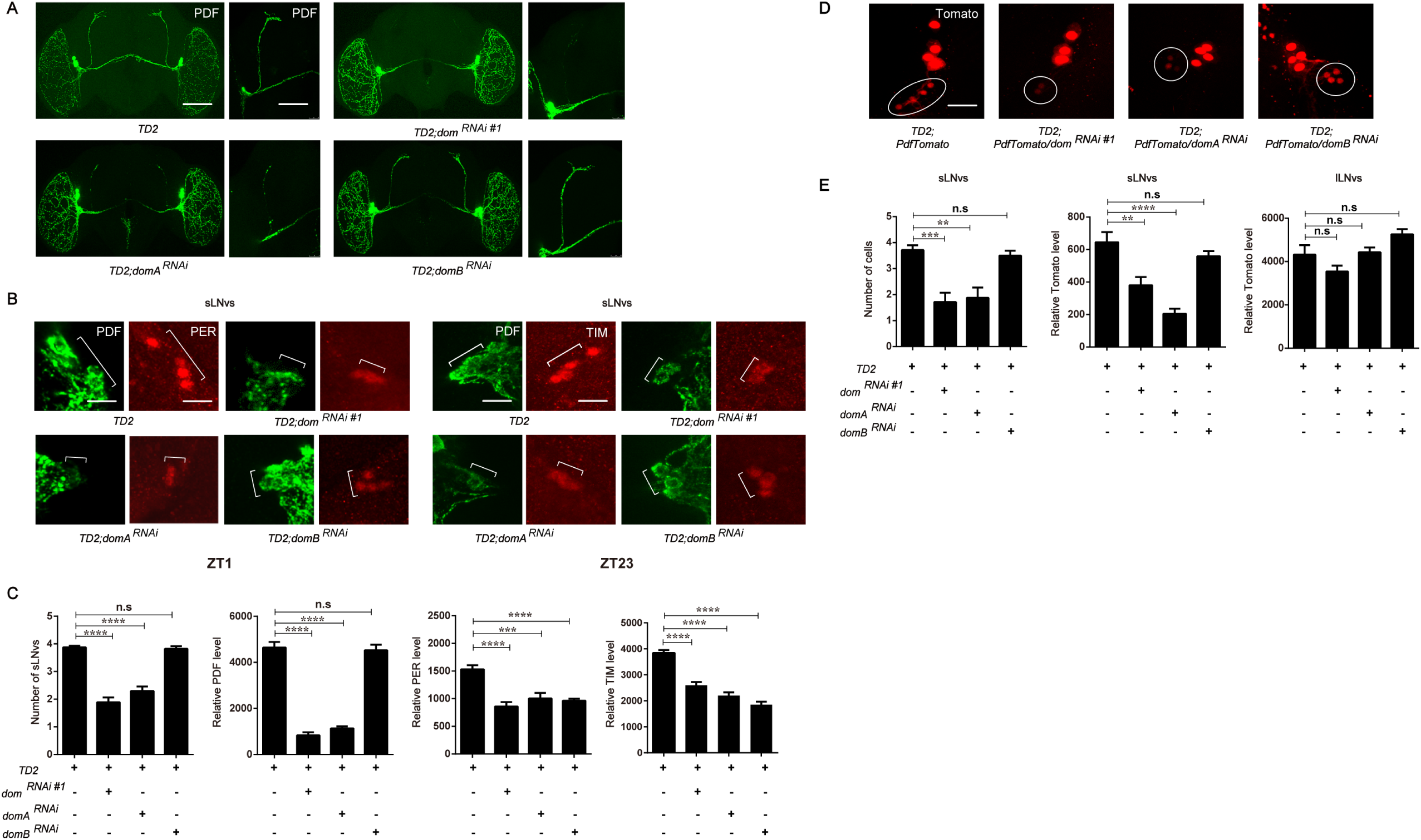
Depletion of DOM-A, but not DOM-B affects s-LN_v_s maintenance and PDF accumulation. **A.** Representative confocal images showing PDF expression in whole brain and dorsal axonal projection of sLN_v_s. Brains were dissected at ZT1 for anti-PDF (green) and anti-PER antibodies (red). Comparing to the control (Top left panel), dorsal projection of sLNvs was disrupted in *dom* (Top right panel) and *domA* downregulation (Bottom left panel) no effects were observed in the presence of *domB* RNAi (Bottom right panel) (Scale bar: left, 500 um; right, 100 um). **B.** Representative images of sLN_v_s of associated lines. Brains were dissected at ZT1 for anti-PDF (green) and anti-PER antibodies (red) or dissected at ZT23 for anti-PDF (green) and anti-TIM antibodies (red). Cell number of sLN_v_s (marked as square brackets), PDF, PER and TIM levels in sLN_v_s were decreased in *dom* and *domA* downregulation flies brains (Top right and bottom left panels), while only affected PER and TIM levels in *domB* downregulation flies (Bottom right panel). (Scale bar: 50 um). **C.** Quantification of sLN_v_s numbers in each brain hemisphere, as well as PDF, PER and TIM levels in sLN_v_s. n=32 hemispheres was used for quantification. **D.** *Pdf* transcriptional level in lLN_v_s and sLN_v_s using TOMATO fluorescence signal. *Pdf* transcriptional levels in sLNvs (marked as circle) were decreased in *dom* and *domA* downregulation fly brains (middle two panels), while sLNv numbers and *pdf* transcriptional levels in *domB* downregulation flies were normal (right panel) (Scale bar: 100 um.). **E.** Quantification of sLN_v_s numbers and TOMATO fluorescence signal in sLN_v_s and lLN_v_s. n=24 hemispheres were used. Error bars correspond to SEM. **p < 0.01,***p < 0.001,****p < 0.0001 as determined by t test.

Next we examined which step of regulation causes the reduction of PDF levels in DOM and DOM-A knockdown. DOM regulates gene expression at the transcription level [22]. We therefore examined the expression of *pdf* using a transcriptional reporter line, *pdf*Tomato [20]. Consistent with PDF staining, we observed significant reduction of *pdf* transcription in sLN_v_s of both DOM and DOM-A knockdown (Fig. 4D, 4E). Interestingly, the decrease in neuron number and reduction of PDF level were unique to sLN_v_s, which were not detected in the lLN_v_s (Fig. 4D, 4E).

In summary, consistent with differences in circadian behavior, DOM-A and DOM-B knockdown also exhibit differences at the molecular level. Knockdown of DOM-A causes loss of sLN_v_s and decrease in sLN_v_ PDF levels, which is not observed in DOM-B downregulation.

### DOM is required during development and adulthood for circadian rhythms

Decrease of sLN_v_s numbers and dorsal projections in adult flies might be due to abnormal development or maintenance defects. Therefore, we performed brain dissections at larval stage and pupal stage. The number of sLN_v_s precursor in the DOM knockdown condition was unchanged compared to the control at 3^rd^ instar larvae (Fig. 5A, 5B), which suggests that DOM does not affect sLN_v_s development. However, same as adulthood, even at early pupal stage (3 days after pupation), there were only 1-2 sLN_v_s detected in *dom* downregulation (Fig. 5C, 5D). These data indicate that DOM is required for the maintenance but not for the development of sLN_v_s. DOM probably affects sLN_v_s during larval-pupal metamorphosis. Consistent with adult flies, both PDF and PER levels of *dom* downregulation were reduced in sLN_v_s compared to the control in larvae or pupae (Fig. 5B, 5D).

**Figure 5.**
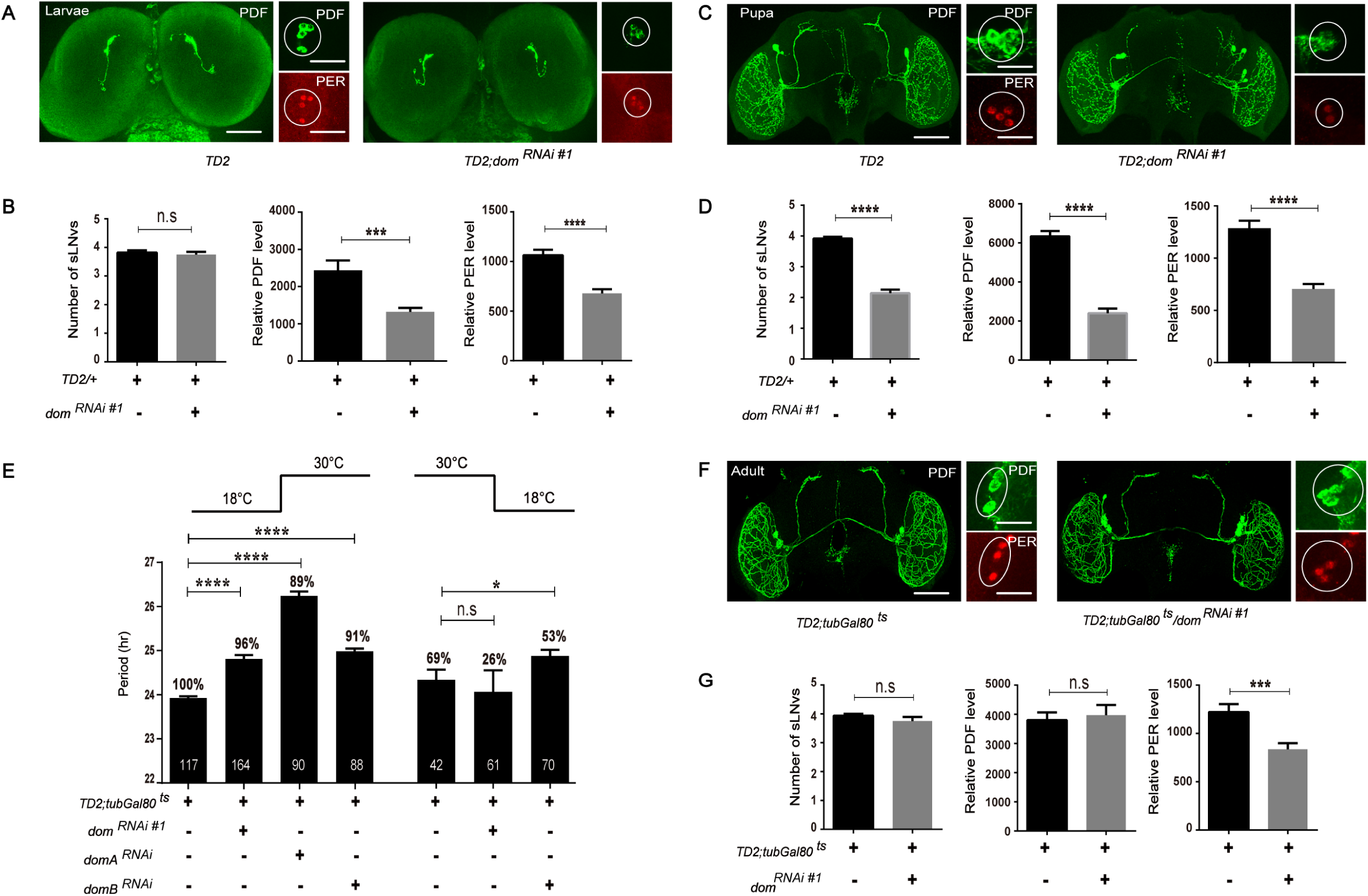
DOM-A is required during development for circadian rhythmicity and adulthood for regulation of period length, while DOM-B is only required in adulthood. **A.** Representative larva brain confocal images. L3 stage larvae collected at ZT1 were immunostained with PDF and PER antibodies. Silence the expression of *dom* in larvae stage affected PDF and PER levels in sLNvs (marked as circle), while number of sLNvs were not affected (Scale bar: left, 200 um; right, 50 um.). **B.** Quantification of larva sLN_v_s, as well as PDF and PER levels. For each genotype, ∼ 22 flies brains and ∼85 neurons were used for quantification. **C.** Representative pupal brain confocal images. Flies pupa (3 days after pupation) were collected at ZT1 were immunostained with PDF and PER antibodies. Silence the expression of *dom* in pupal stage affected number of sLNvs, PDF and PER levels in sLNvs (marked as circle) (Scale bar: left, 500 um; right, 50 um). **D.** Quantification of pupal sLN_v_s, as well as PDF and PER levels. For each genotype, ∼ 25 flies brains and ∼100 neurons were used for quantification. **E.** Free-running period and percentage of rhythmicity of DOM, DOM-A and DOM-B depleted flies in adulthood (left part) or during development (right). Stage specific silencing the was done by using the conditional *tim-Gal4;tub-Gal80*^*ts*^ driver system. **F**. Representative confocal images of fly brain showing projections of PDF positive LN_v_s. Flies were grown at 18°C until eclosion, and adult flies were entrained for 5 days in LD 30°C. Brains were dissected at ZT1 for anti-PDF (green) and anti-PER antibodies (red) (Scale bar: left, 500 um; right, 50 um). **G**. Quantification of the numbers of sLNvs and PDF, PER levels in sLN_v_s of associated lines. Error bars correspond to SEM. \**P* < 0.05,***p < 0.001, ****p < 0.0001 as determined by the *t*-test.

The decrease of axonal projections and number of sLN_v_s we observed in DOM knockdown flies (Fig. 4A, 4B) suggested that DOM might play a role during development to properly regulate circadian rhythms. Thus, we used *tim-GAL4* and GAL80^ts^ TARGET system to temporally express *dom* dsRNA during development or after eclosion [40]. 18 °C prevents dsRNA production, while 30°C allows expression of dsRNA thus downregulation of DOM. We observed strong rhythmicity but around 1-2 hr period-lengthening with adult-specific DOM or DOM-A knockdown, compared to high arrhythmic activity for knockdown during development (Fig. 5E and Appendix Table S1). Thus, developmental expression of *dom* or *dom-A* dsRNA appears to affect the rhythmicity, while adult specific depletion appears to be sufficient for period-lengthening.

Furthermore, knockdown of DOM-B only in adulthood caused a ∼1 hr increase in period length, suggesting that unlike DOM-A, the DOM-B splice form is mainly necessary for the regulation of circadian clocks in adult stage (Fig. 5E and Appendix Table S1). We observed a slight but significant lengthening of period for DOM-B developmental depletion, which indicates that DOM-B might also play a role during development.

To exclude anatomical defects in the circadian circuitry, we performed brain dissections in the flies with adult-specific DOM downregulation. As expected, the dorsal projections and number of sLN_v_s were normal, and PDF level was not affected (Fig. 5F, 5G). These data further suggest that the arrhythmia we observed in DOM knockdown was mainly due to the function of DOM during development. However, the abundance of PER in sLN_v_s was still reduced, which could explain the lengthened period phenotype (Fig. 5G). Taken together, these data indicate that DOM-A is required during development for the rhythmic activity, while both DOM-A and DOM-B are necessary for period determination post development.

### Activation of PDFR signaling in circadian neurons rescues the behavioral phenotype of DOM depletion

Since we observed low PDF levels in the sLN_v_s with DOM knockdown, we wondered whether the decrease of PDF signaling was the cause of arrhythmia. To test this hypothesis, we determined whether hyperactivation of PDFR could rescue the phenotypes that accompany reduction in DOM expression. To increase PDFR signaling, we expressed a membrane-tethered PDF (t-PDF), which has been shown to mimic high PDF levels [41]. A scrambled peptide sequence (t-PDF SCR) with similar length as PDF was used as a negative control. Strikingly, coexpression of t-PDF restored the rhythmicity to 89% for *dom* RNAi (71% for another independent *dom* RNAi line, Fig. 6A, 6B), whereas the t-PDF SCR control was unable to restore arrhythmicity (Fig. 6A, 6B). Interestingly, not only the rhythmicity, but also the period of DOM knockdown was rescued with coexpression of t-PDF (Fig. 6B). We then stained fly brains with PDF and pacemaker proteins PER and TIM (Fig. 6C and Appendix Fig S6). With expression of t-PDF in DOM knockdown flies, we made two interesting observations. First, PDF level as well as the dorsal projection of sLN_v_s were restored, while the number of sLN_v_s was still lower (Fig. 6C, 6D). Second, the abundance of PER and TIM in sLN_v_s was also rescued (Fig. 6C, 6D and Appendix Fig S6). This might be due to the fact that PDF stabilizes PER and TIM [16,17].

**Figure 6.**
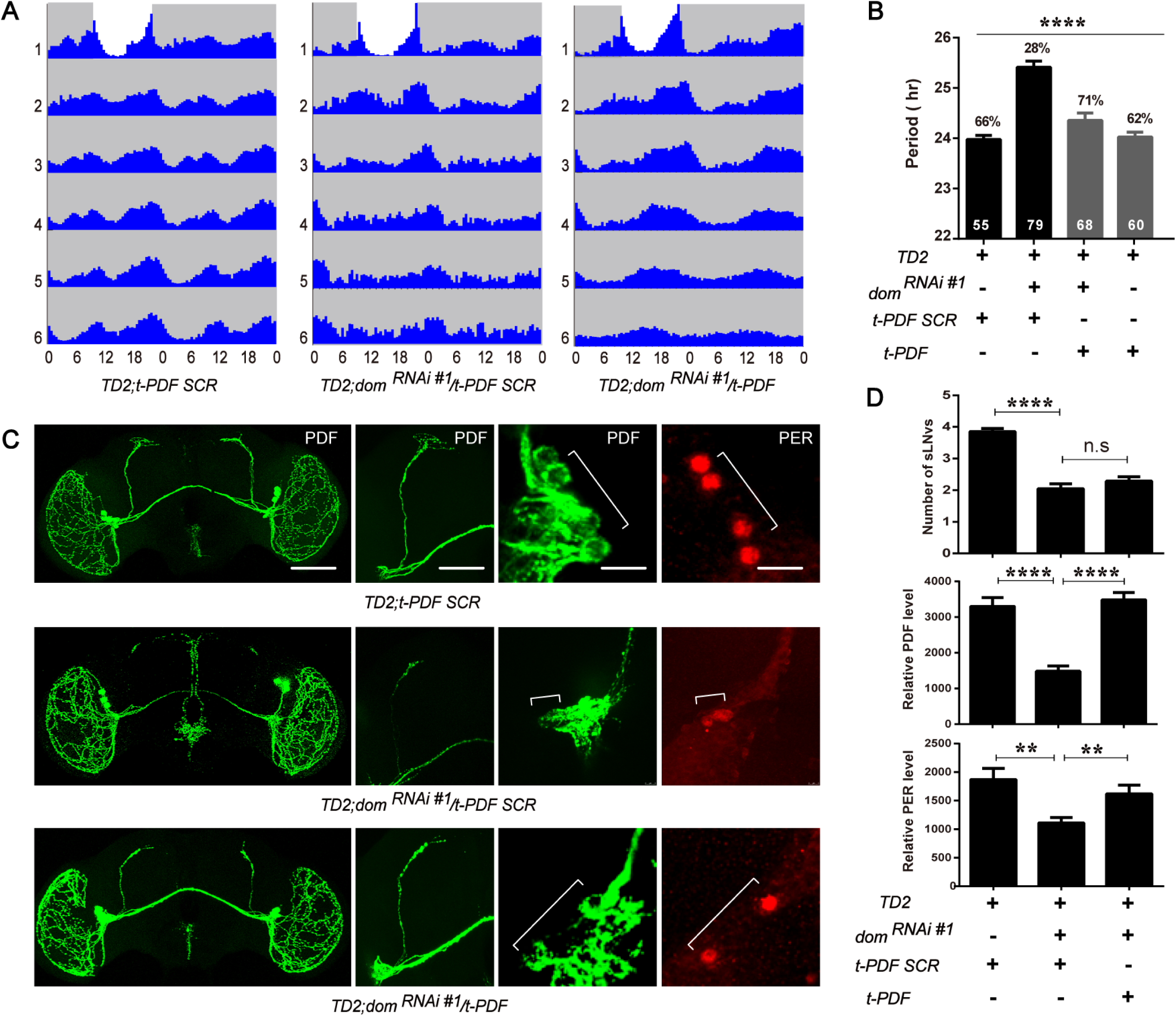
Constitutive activation of PDFR signaling rescued the circadian behavior phenotype of DOM downregulation. **A.** Actograms showing the average activities on the last day of LD and during 5 days in DD. Light represents the day and gray darkness. From left to right: (Left panel) flies expressing the membrane-tethered scrambled PDF; (middle panel) *dom*^*RNAi#1*^ flies expressing a membrane-tethered scrambled PDF (negative control); and (right panel) *dom*^*RNAi#1*^ flies expressing the membrane-tethered PDF. Arrhythmia and long period of *dom* downregulation are rescued with tethered PDF. **B**. Free-running period of *dom* RNAi flies expressing the membrane-tethered PDF. **C**. Representative confocal images of brains of *dom* RNAi flies expressing the membrane-tethered PDF or scrambled PDF. Flies were entrained for 4 days in LD 25°C, and brains were dissected at ZT1 for anti-PDF antibody (green) and anti-PER antibody (red). From top to bottom: (Top panel) fly brain expressing the membrane-tethered scrambled PDF; (middle panel) *dom*^*RNAi#1*^ flies expressing a membrane-tethered scrambled PDF; and (bottom panel) *dom*^*RNAi#1*^ flies expressing the membrane-tethered PDF. Confocal images are whole brain, dorsal projection and soma of sLN_v_s from left to the right (Scale bar: whole brain, 500 um; projection, 100 um; sLNvs, 50 um). **D**. Quantification of the number and relative PDF and PER levels of sLNvs. For each genotype, totally, 20-25 flies brains and 50-80 neurons were used for quantification of the staining. Error bars correspond to SEM. **p < 0.01, ****p < 0.0001 as determined by *t*-test.

## Discussion

Here, we identify that two alternatively spliced variants of DOM play distinct roles in circadian rhythms. DOM-A is critical for maintenance of sLN_v_s and PDF abundance during development thus controls morning anticipatory activity and circadian rhythmicity; while DOM-B specifically determines circadian period of locomotor activity by regulating core pacemaker protein PER/TIM abundance.

The abolished morning anticipatory activity in light dark cycle and low rhythmicity under constant darkness in DOM/DOM-A knockdown flies is reminiscent of the phenotypes seen in *pdf* mutant (Fig.1 and Fig.3) [42]. Based on these observations, we hypothesized that PDF signaling is disrupted with *dom* or *dom-A* RNAi. Indeed, we found that PDF abundance was decreased both in the soma and dorsal axonal projections of sLN_v_s (Fig. 4). Furthermore, by restricting RNAi to adulthood, we rescued the PDF expression and projection, which also restored the rhythms (Fig. 5). These results suggest that the arrhythmic phenotype is due to the decrease of PDF levels in *dom* or *dom-A* downregulation. Despite the importance of sLN_v_s in circadian rhythms, the mechanism underlying sLN_v_s maintenance is still unclear. Here we find that DOM-A plays important roles in the maintenance of sLN_v_s. PDF positive sLN_v_s and dorsal projections are formed in first instar larvae [10]. During development, both DOM-A and DOM-B start to be expressed in embryos [23]. Our data suggest that DOM-A does not affect sLN_v_s development in larvae, but rather regulates sLN_v_s maintenance at later stages (Fig. 5). Knockdown of DOM-A may trigger apoptosis or necrosis pathways and cause programmed cell death in sLN_v_s. In fact, chromatin-modifying pathway has been shown to regulate neuronal necrosis in flies [43]. In the future, it will be interesting to examine whether DOM-A is involved these programmed cell death pathways and blocking necrosis or apoptosis pathway can restore the sLN_v_s in DOM downregulation.

Here we found that depletion of DOM-A seems to specifically affect the maintenance of PDF positive sLN_v_s and PDF accumulation in sLN_v_s, while the PDF positive lLN_v_s is not affected (Fig. 4). This suggests that as master pacemaker neurons, sLN_v_s may have some unique regulatory program for maintenance and gene expression, which requires DOM-A. Interestingly, a phosphatase LAR has recently been found to specifically regulate PDF expression in the dorsal projection of sLN_v_s during development [39]. The pacemaker protein VRI also played important roles in controlling PDF abundance and dorsal arborization rhythms of sLN_v_s [21]. Future efforts to identify specific regulatory mechanisms in sLN_v_s will help us understand the maintenance and function of circadian neuronal network.

Why does DOM-A, but not DOM-B, regulate sLN_v_ maintenance and *pdf* transcription? One possibility might be that DOM-A and DOM-B are associated with different protein complexes. Compared to DOM-B, DOM-A has two unique domains in the C-terminus: a SANT domain and a poly-Q domain. Both of these two domains are known to mediate protein interactions. Interestingly, only DOM-A is identified to bind the Tip60 complex when Tip60 was purified by a tagged protein subunit from *Drosophila* S2 cells [22]. A previous study finds that depletion of Tip60 or overexpression of a histone acetyltransferase-defective Tip60 decreases axonal growth of the sLN_v_s in the fly model of Alzheimer’s disease [44]. These results suggest that DOM-A might interact with Tip60 to control the axonal projection and PDF expression of sLN_v_s.

Lastly, how does DOM control the length of circadian period? This is probably through regulation of PER and TIM abundance. Interestingly, compared to the decrease of *per* and *tim* mRNA in DOM downregulation, the master circadian transcription factor *clk* mRNA is not affected. Does DOM control PER and TIM accumulation through transcriptional regulation? Indeed, we found that from *Drosophila* S2 cell affinity pull-down, CLK interacts with DOM complex (Table 1). Furthermore, DOM-A and DOM-B binding is enriched in the E-box region of *per* and *tim* promoters. Even though no binding differences were observed in DOM-A and DOM-B at ZT4 and ZT16 (Fig. 3), it is possible that other subunits in the DOM complex bind rhythmically, leading to rhythmic activity of the complex. Given that the conserved functions of DOM and human SRCAP/p400 in Notch signaling and histone variant exchange, it is possible that similar mechanisms are leveraged to control circadian rhythms in mammals.

## Materials and Methods

### Fly Stocks

All the flies were raised on cornmeal/agar medium at 25°C under a LD cycle. The following strains were used: *yw, w*^*1118*^, *yw; tim-GAL4/CyO* [35], *yw;Pdf-GAL4/CyO* [11]; *yw;tim-GAL4/CyO;Pdf-GAL80/TM6B* [45], *tublin-GAL80*^*ts*^, *UAS-cd8-GFP, clk*^*out*^. The following stocks were obtained from the Bloomington Drosophila Stock Center (http://flystocks.bio.indiana.edu/): *UAS-dom*^*RNAi#1*^ (BL31054), *UAS-dom*^*RNAi#2*^ (BL38941), *UAS-domA* (BL64261) and *UAS-domB* (BL64263). *DomA*^*RNAi*^ *(sh-domA)* and *domB*^*RNAi*^ *(sh-domB)* fly lines were generous gifts from Dr. Peter B. Becker. *PdfTomato* line was generated by Dr. Sebastian Kadener.

### Behavioral Experiments and Analysis

For behavioral experiments, adult male flies (2-4 days old) were used for testing locomotor activity rhythms. Flies were entrained for 4 full days LD cycle at 25°C, using about 500 lux light intensities, and then released into DD at 25°C for at least 6 days. Locomotor activity was measured with TriKinetics Activity Monitors in I36-LL Percival Incubators. Locomotor activity was averaged over the 4 days entrainment for LD and 6 days for DD. Analyses of period and power were carried out using FaasX software as previously described [46]. Actograms were generated using a signal-processing toolbox implemented in MATLAB (MathWorks) [47]. For *GAL80*^*ts*^ experiments, flies were raised at 18°C and tested at 30°C. They were entrained for 5 days and then released in DD for at least 6 days. Morning anticipation was calculated by the ratio of activity counts between 2 hrs before light on and 6 hrs before light on. We first measured the single fly activity counts obtained in twelve 30-min bins between Zeitgeber Time (ZT) 17.5 and ZT24 (6 hr before lights on) and six 30-min bins between ZT22.5 and ZT24 (2 hrs before lights on). The first value is divided by the second to obtain the morning activity index. Morning anticipations of individual flies were then averaged and plotted on the graphs.

### Whole-Mount Immunohistochemistry

Whole-mount immunohistochemistry for fly brains were done as previously described [48]. Adult fly (3-6 days old) or L3 stage larval brains were dissected in chilled PBT (PBS with 0.1% Triton X-100) at the indicated time points and fixed in 4% formaldehyde diluted in PBS for 30 min at room temperature. For pupal brains dissection, *Drosophila* were fully developed and hatched 7 days after pupation in the same cross, pupa were collected 3 days after pupation using a small wet paintbrush and transfer to a dissection dish. After 2-3 times PBS wash, the pupal brains were dissected and fixed as previously described method. The brains were rinsed and washed with PBT three times (10 min each). Then, brains were incubated with 10% normal Goat serum diluted in PBT to block for 60 min at room temperature and incubated with primary antibodies at 4°C overnight. For PER, TIM and CLK staining, we used 1:1,500 rabbit anti-PER, 1:2,000 rat anti-TIM (gift from Dr. Rosbash) and guinea pig 1:2,500 anti-CLK (gift from Dr. Hardin), respectively. We used a 1:200 dilutions for mouse anti-PDF and 1:200 for rabbit anti-GFP (DSHB). After six washes with PBT (20 min each), brains were incubated with relative secondary antibody at 4°C overnight, followed by another six washes with PBT. For *PdfTomato* staining, flies were directly dissected in chilled PBT at the indicated time points and moutained in the medium (VECTOR). All samples were imaged on a Leica Confocal SP8 system, with laser settings kept constant within each experiment. 10 to 12 fly brains for each genotype were dissected for imaging. Representative images are shown. Image-J software (National Institutes of Health [NIH]) was used for PER, PDF, TIM and CLK quantification in 15-30 sLNvs from at least seven brains. For quantification, signal intensity in each sLNv and average signals in three neighboring non-circadian neurons were measured, and the ratio between signals in sLNvs and non-circadian neurons was calculated.

### Chromatin immunoprecipitation (ChIP)

Chromatin immunoprecipitation (ChIP) was performed based on published protocols [49]. Flies entrained in 12 hr light:12 hr dark (LD) conditions at 25°C for four days were collected at two time-points (ZT) on the fifth day. Briefly, chromatin was isolated from 500 μl of fly heads homogenized with a dounce homogenizer (Kimble Chase) for 20 strokes using the loose “A” pestle. Homogenate was sieved by a 70 μm cell strainer (Falcon) then centrifuged to remove cell debris. Pellets were cross-linked using formaldehyde. Samples were sonicated using a focused-ultrasonicator (Covaris M220) on setting for 400-500 bp cDNA and then centrifuged at 10,000 rpm for 10 minutes. Supernatant was collected in two 130 μl aliquots for IP and 26 μl was collected for input and frozen at −80C for analysis. Sonicated chromatin was roughly 500 bp in length (<1000 bp). For each IP, 30 μl of a Protein G Dynabead slurry (Life Technologies) was washed then incubated along with the appropriate antibodies for 4 hours at 4°C with rotation. Amount of antibodies used for ChIP is as follows: anti-DOM-A (20 μg/ml, rabbit, GenScript) which was generated in our lab (antigen protein sequence designed according to 2008-2349 aa of DOM-A), anti-rabbit-IgG (20 μg/ml, Life Technologies), anti-DOM-B (10 μg/ml, mouse, from Dr. Peter B. Becker), anti-mouse-IgG (10 μg/ml, Life Technologies). Following incubation, beads were collected and incubated with chromatin overnight at 4°C with rotation. DNA was eluted using the Qiagen PCR purification kit and subjected to qPCR. At least three technical replicates of qPCR were performed for each biological ChIP replicate and three biological replicates were performed for DOM-A and DOM-B assays. Background binding to a nonspecific antibody (anti-IgG; Life Technologies) bound to Dynabeads was subtracted from input samples and results are presented as the percentage of the input samples. For each assay, at least three biological replicates were performed. The specific primers used for qPCR are described in Appendix Table S2. The technical qPCR triplicates were averaged for each biological replicate as no significant differences were found between the technical replicates, and the error bars represent SEM calculated from variance between biological replicates. Two-tailed t-tests were used to determine statistical differences between control and experimental treatment at each ZT.

### Real time quantitative reverse transcription PCR

Flies were collected at the indicated time points and isolated heads were stored at −80°C. Total RNA was extracted from 25–30 heads with TRIzol based on the manufactures protocol (Life Technologies, USA). A 2-µg quantity of RNA was reverse-transcribed with reverse transcription reagents (Invitrogen). For real-time PCR (qRT-PCR) of *per*, *tim*, *clk*, *dom*, *domA*, *domB* and *actin*, we used a qPCR detection kit (SYBR Select Master Mix For CFX) (Life Technologies, USA). The specific primers used for qPCR are described in Appendix Table S2. All the experiments were performed in the CFX96 Real-Time System (BIO-RAD).

### Western blot analysis

Fly heads were collected at the indicated time points and homogenized with pestles, protein extracts were prepared with HEPES-Triton lysis buffer (1X HEPES-Triton buffer, 1 mM DTT, 0.4%NP-40, 0.1% SDS, 10% glycerol, 1X tablet protease inhibitor). Proteins were quantified using BSA assay. For immunoblot analysis, proteins were transferred to PVDF membranes (Genesee Scientific) and incubated with anti-DOM-A (1:300, GenScript), anti-DOM-B (1:5, from Dr. Peter B. Becker) and anti-ACTIN (1:100,DSHB) in blocking solution. Band intensity was calculated and analysed with Image J.

### Statistics analysis

Statistical analysis of two data points was performed with Student’s t-test. Statistical analysis of multiple data points was performed with one-way analysis of variance with Tukey post-hoc tests using GraphPad software.

## Acknowledgements

We would like to thank Drs. Luoying Zhang, Patrick Emery, Pedro Miura, and Alexander M. van der Linden for comments and discussions on this manuscript. We thank all the members in the Zhang lab for technical support and helpful discussions. We thank Dr. Becker for the fly strains of *domA* and *domB* shRNA, as well as the DOM antibodies. We thank the Bloomington stock center for various fly stocks. We are grateful for TIM and PER antibodies kindly given by Dr. Rosbash and Dr. Stanewsky. We also thank the Developmental Studies Hybridoma Bank for PDF and GFP antibodies. This work was initiated in Dr. Patrick Emery’s lab. This work is supported by the National Institute of General Medical Sciences of the National Institutes of Health under grant numbers P20 GM103650, GM103554, and GM103440, and R01 GM102225 to JCC.

## Author Contributions

YZ, JCC and ZL formulated the ideas and designed the experiments. ZL and VHL performed the experiments. YZ, JCC and ZL wrote the manuscript.

## References

1. Bell-Pedersen D, Cassone VM, Earnest DJ, Golden SS, Hardin PE, Thomas TL, Zoran MJ (2005) Circadian rhythms from multiple oscillators: Lessons from diverse organisms. Nature Reviews Genetics 6: 544–556

2. Hardin PE, Panda S (2013) Circadian timekeeping and output mechanisms in animals. Curr Opin Neurobiol 23: 724–731

3. Tataroglu O, Emery P (2015) The molecular ticks of the Drosophila circadian clock. Current Opinion in Insect Science 7: 51–57

4. Nitabach MN, Taghert PH (2008) Organization of the Drosophila circadian control circuit. Curr Biol 18: R84–93

5. Grima B, Chelot E, Xia R, Rouyer F (2004) Morning and evening peaks of activity rely on different clock neurons of the Drosophila brain. Nature 431: 869–873

6. Stoleru D, Peng Y, Agosto J, Rosbash M (2004) Coupled oscillators control morning and evening locomotor behaviour of Drosophila. Nature 431: 862–868

7. Gorostiza EA, Depetris-Chauvin A, Frenkel L, Pirez N, Ceriani MF (2014) Circadian pacemaker neurons change synaptic contacts across the day. Curr Biol 24: 2161– 2167

8. Sivachenko A, Li Y, Abruzzi KC, Rosbash M (2013) The transcription factor Mef2 links the Drosophila core clock to Fas2, neuronal morphology, and circadian behavior. Neuron 79: 281–292

9. Depetris-Chauvin A, Fernandez-Gamba A, Gorostiza EA, Herrero A, Castano EM, Ceriani MF (2014) Mmp1 Processing of the PDF Neuropeptide Regulates Circadian Structural Plasticity of Pacemaker Neurons. Plos Genetics 10:

10. HelfrichForster C (1997) Development of pigment-dispersing hormone-immunoreactive neurons in the nervous system of Drosophila melanogaster. Journal of Comparative Neurology 380: 335–354

11. Renn SCP, Park JH, Rosbash M, Hall JC, Taghert PH (1999) A pdf neuropeptide gene mutation and ablation of PDF neurons each cause severe abnormalities of behavioral circadian rhythms in Drosophila. Cell 99: 791–802

12. Yao Z, Shafer OT (2014) The Drosophila Circadian Clock Is a Variably Coupled Network of Multiple Peptidergic Units. Science 343: 1516–1520

13. Mertens I, Vandingenen A, Johnson EC, Shafer OT, Li W, Trigg JS, De Loof A, Schoofs L, Taghert PH (2005) PDF receptor signaling in Drosophila contributes to both circadian and geotactic behaviors. Neuron 48: 213–219

14. Lear BC, Lin JM, Keath JR, McGill JJ, Raman IM, Allada R (2005) The ion channel narrow abdomen is critical for neural output of the Drosophila circadian pacemaker. Neuron 48: 965–976

15. Hyun S, Lee Y, Hong ST, Bang S, Paik D, Kang JK, Shin J, Lee J, Jeon K, Hwang S, et al. (2005) Drosophila GPCR Han is a receptor for the circadian clock neuropeptide PDF. Neuron 48: 267–278

16. Li Y, Guo F, Shen J, Rosbash M (2014) PDF and cAMP enhance PER stability in Drosophila clock neurons. Proceedings of the National Academy of Sciences of the United States of America 111: E1284–E1290

17. Seluzicki A, Flourakis M, Kula-Eversole E, Zhang LY, Kilman V, Allada R (2014) Dual PDF Signaling Pathways Reset Clocks Via TIMELESS and Acutely Excite Target Neurons to Control Circadian Behavior. Plos Biology 12:

18. Blau J, Young MW (1999) Cycling vrille expression is required for a functional Drosophila clock. Cell 99: 661–671

19. Park JH, Helfrich-Forster C, Lee G, Liu L, Rosbash M, Hall JC (2000) Differential regulation of circadian pacemaker output by separate clock genes in Drosophila. Proc Natl Acad Sci U S A 97: 3608–3613

20. Mezan S, Feuz JD, Deplancke B, Kadener S (2016) PDF Signaling Is an Integral Part of the Drosophila Circadian Molecular Oscillator. Cell Rep 17: 708–719

21. Gunawardhana KL, Hardin PE (2017) VRILLE Controls PDF Neuropeptide Accumulation and Arborization Rhythms in Small Ventrolateral Neurons to Drive Rhythmic Behavior in Drosophila. Curr Biol 27: 3442–3453 e3444

22. Kusch T, Florens L, MacDonald WH, Swanson SK, Glaser RL, Yates JR, Abmayr SM, Washburn MP, Workman JL (2004) Acetylation by Tip60 is required for selective histone variant exchange at DNA lesions. Science 306: 2084–2087

23. Ruhf ML, Braun A, Papoulas O, Tamkun JW, Randsholt N, Meister M (2001) The domino gene of Drosophila encodes novel members of the SW12/SNF2 family of DNA-dependent ATPases, which contribute to the silencing of homeotic genes. Development 128: 1429–1441

24. Borner K, Becker PB (2016) Splice variants of the SWR1-type nucleosome remodeling factor Domino have distinct functions during Drosophila melanogaster oogenesis. Development 143: 3154–3167

25. Tea JS, Luo LQ (2011) The chromatin remodeling factor Bap55 functions through the TIP60 complex to regulate olfactory projection neuron dendrite targeting. Neural Development 6:

26. Rust K, Tiwari MD, Mishra VK, Grawe F, Wodarz A (2018) Myc and the Tip60 chromatin remodeling complex control neuroblast maintenance and polarity in Drosophila. Embo J, 10.15252/embj.201798659

27. Dubruille R, Murad A, Rosbash M, Emery P (2009) A Constant Light-Genetic Screen Identifies KISMET as a Regulator of Circadian Photoresponses. Plos Genetics 5:

28. Kwok RS, Lam VH, Chiu JC (2015) Understanding the role of chromatin remodeling in the regulation of circadian transcription in Drosophila. Fly 9: 145–154

29. Adewoye AB, Kyriacou CP, Tauber E (2015) Identification and functional analysis of early gene expression induced by circadian light-resetting in Drosophila. Bmc Genomics 16:

30. Kwok RS, Lam VH, Chiu JC (2015) Understanding the role of chromatin remodeling in the regulation of circadian transcription in Drosophila. Fly 9: 145–154

31. Choi H, Larsen B, Lin ZY, Breitkreutz A, Mellacheruvu D, Fermin D, Qin ZS, Tyers M, Gingras AC, Nesvizhskii AI (2011) SAINT: probabilistic scoring of affinity purification-mass spectrometry data. Nature methods 8: 70–73

32. Kula-Eversole E, Nagoshi E, Shang YH, Rodriguez J, Allada R, Rosbash M (2010) Surprising gene expression patterns within and between PDF-containing circadian neurons in Drosophila. P Natl Acad Sci USA 107: 13497–13502

33. Dietzl G, Chen D, Schnorrer F, Su KC, Barinova Y, Fellner M, Gasser B, Kinsey K, Oppel S, Scheiblauer S, et al. (2007) A genome-wide transgenic RNAi library for conditional gene inactivation in Drosophila. Nature 448: 151–U151

34. Zhang XY, Lu K, Zhou JL, Zhou Q (2013) Molecular characterization and gene functional analysis of Dicer-2 gene from Nilaparvata lugens (Hemiptera: Geometroidea). Insect Science 20: 61–68

35. Kaneko M, Hall JC (2000) Neuroanatomy of cells expressing clock genes in Drosophila: Transgenic manipulation of the period and timeless genes to mark the perikarya of circadian pacemaker neurons and their projections. Journal of Comparative Neurology 422: 66–94

36. Mahesh G, Jeong E, Ng FS, Liu Y, Gunawardhana K, Houl JH, Yildirim E, Amunugama R, Jones R, Allen DL, et al. (2014) Phosphorylation of the transcription activator CLOCK regulates progression through a approximately 24-h feedback loop to influence the circadian period in Drosophila. The Journal of biological chemistry 289: 19681–19693

37. Abruzzi KC, Rodriguez J, Menet JS, Desrochers J, Zadina A, Luo W, Tkachev S, Rosbash M (2011) Drosophila CLOCK target gene characterization: implications for circadian tissue-specific gene expression. Genes Dev 25: 2374–2386

38. Veleri S, Rieger D, Helfrich-Forster C, Stanewsky R (2007) Hofbauer-Buchner eyelet affects circadian photosensitivity and coordinates TIM and PER expression in Drosophila clock neurons. Journal of Biological Rhythms 22: 29–42

39. Agrawal P, Hardin PE (2016) The Drosophila Receptor Protein Tyrosine Phosphatase LAR Is Required for Development of Circadian Pacemaker Neuron Processes That Support Rhythmic Activity in Constant Darkness But Not during Light/Dark Cycles. J Neurosci 36: 3860–3870

40. McGuire SE, Roman G, Davis RL (2004) Gene expression systems in Drosophila: a synthesis of time and space. Trends in Genetics 20: 384–391

41. Choi C, Fortin JP, McCarthy EV, Oksman L, Kopin AS, Nitabach MN (2009) Cellular Dissection of Circadian Peptide Signals with Genetically Encoded Membrane-Tethered Ligands. Current Biology 19: 1167–1175

42. Renn SC, Park JH, Rosbash M, Hall JC, Taghert PH (1999) A pdf neuropeptide gene mutation and ablation of PDF neurons each cause severe abnormalities of behavioral circadian rhythms in Drosophila. Cell 99: 791–802

43. Liu K, Ding L, Li Y, Yang H, Zhao C, Lei Y, Han S, Tao W, Miao D, Steller H, et al. (2014) Neuronal necrosis is regulated by a conserved chromatin-modifying cascade. Proc Natl Acad Sci U S A 111: 13960–13965

44. Pirooznia SK, Sarthi J, Johnson AA, Toth MS, Chiu K, Koduri S, Elefant F (2012) Tip60 HAT Activity Mediates APP Induced Lethality and Apoptotic Cell Death in the CNS of a Drosophila Alzheimer’s Disease Model. Plos One 7:

45. Murad A, Emery-Le M, Emery P (2007) A subset of dorsal neurons modulates circadian behavior and light responses in Drosophila. Neuron 53: 689–701

46. Grima B, Lamouroux A, Chelot E, Papin C, Limbourg-Bouchon B, Rouyer F (2002) The F-box protein Slimb controls the levels of clock proteins Period and Timeless. Nature 420: 178–182

47. Levine JD, Funes P, Dowse HB, Hall JC (2002) Signal analysis of behavioral and molecular cycles. BMC Neurosci 3: 1

48. Zhang Y, Ling J, Yuan C, Dubruille R, Emery P (2013) A role for Drosophila ATX2 in activation of PER translation and circadian behavior. Science 340: 879–882

49. Kwok RS, Li YH, Lei AJ, Edery I, Chiu JC (2015) The Catalytic and Non-catalytic Functions of the Brahma Chromatin-Remodeling Protein Collaborate to Fine-Tune Circadian Transcription in Drosophila. PLoS genetics 11: e1005307

